# Novel clonal lineages of wheat stem rust in the Southern Cone of South America

**DOI:** 10.64898/2025.12.18.695303

**Authors:** Camilla Langlands-Perry, Eva C. Henningsen, Clare M. Lewis, Oadi Matny, Rebecca Spanner, Brian J. Steffenson, Jana Sperschneider, Silvia German, Diane G.O. Saunders, Peter N. Dodds, Pablo Campos, Paula Silva, Melania Figueroa

## Abstract

*Puccinia graminis* f. sp. *tritici* (*Pgt*) causes wheat stem rust, a devastating disease of cereals. Recent approaches to examine populations at a genomic level have provided valuable information on the genotypic diversity of *Pgt* populations in Africa, Europe and North America and evolutionary mechanisms underlying the emergence of new races. However, an in-depth characterisation of *Pgt* populations in South America has been lacking. To bridge this knowledge gap, 91 *Pgt* isolates were collected from Argentina and Uruguay in 2020 and 2021 and used to generate transcriptome and whole genome sequence data and pathotype information. Phylogenetic analyses revealed that this South American *Pgt* population includes three clonal lineages, two of which have not been detected elsewhere. The third lineage is globally dispersed, including isolates from Africa and the Middle East. The predominant lineage, unique to South America, encompassed 90% of the samples and showed related pathotypes differing by virulence on single resistance genes consistent with evolution by stepwise mutation within the clonal lineage. There was no evidence of sexual recombination giving rise to new genetic diversity in this population. These observations urge to routinely incorporate and compare genotypic data from South American *Pgt* isolates to surveillance data from other geographic regions.

## Manuscript body

Wheat stem rust, caused by *Puccinia graminis* f. sp. *tritici* (*Pgt*), is a major global disease of wheat and other cereals (rye, barley and triticale), which has experienced an alarming resurgence over the last 25 years after having previously been regarded as a dormant threat (Singh et al. 2015; Saunders et al. 2019). The emergence of novel races of *Pgt*, with new virulence combinations has resulted in significant epidemics such as those caused by the notorious Ug99 lineage, TTKSK and derived races (Pretorius et al. 2000). Additional new lineages represented by the TKTTF and TTRTF races have caused other epidemics in Africa and Europe (Olivera et al. 2015; Bhattacharya 2017). These outbreaks highlight the vulnerability of wheat production and the importance of maintaining global surveillance networks.

Strategies to prevent or control rust disease are generally based on the introduction of resistance (*R*) genes into cultivars through conventional breeding (Periyannan et al. 2017). These *R* genes often encode receptors that can activate an immune response upon recognition of a corresponding avirulence (Avr) effector protein from the pathogen (Ngou et al. 2022; Dodds 2023). Resistance conferred by *R* genes imposes high selection pressure on pathogen populations favouring mutations in *Avr* genes to escape recognition, which can limit resistance durability (Figueroa et al. 2020). Thus, new races of *Pgt* can arise by single-step mutation of existing clonal lineages to overcome specific *R* genes. Alternatively, new genetic lineages with novel virulence combinations may arise by genetic recombination as a result of sexual reproduction or through nuclear exchange by somatic hybridisation between existing clonal lineages (Figueroa et al. 2020; Henningsen et al. 2025). The latter process is unique to dikaryotic fungi like the rusts, which contain two distinct haploid nuclei per cell. For example, the Ug99 lineage (clade I) emerged by a somatic hybridization event between an isolate of the widespread Pgt21 lineage (clade IX) and another of an African lineage (clade II), which each donated one haploid nucleus to form the new Ug99 lineage (Li et al. 2019; Henningsen et al. 2025). Importantly, sexual recombination is limited by the availability of an alternate host (barberry or mahonia) that is required to allow progression of *Pgt* through this stage of its lifecycle. Barberry and mahonia absent or rare in many wheat-growing regions, restricting the pathogen to asexual reproduction. However, local sexual populations occur in some regions where these hosts are present, which could contribute to the emergence of new strains (Saunders et al. 2019; Olivera et al. 2022; Tsushima et al. 2022; Rodriguez-Algaba et al. 2023). For instance, the origin of the TKTTF race is believed to have been driven by sexual reproduction (Olivera Firpo et al. 2017; Olivera et al. 2019).

The resurgence of stem rust in Africa and Europe has resulted in close monitoring of *Pgt* populations in these regions through genomics-driven analysis of genetic diversity. A SNPchip genotyping platform with about ∼3,000 SNPs (Olivera Firpo et al. 2017) was initially used to define several stem rust clonal lineages in Europe and Africa, while further analysis of *Pgt* isolates by transcriptome or whole genome sequencing has identified a number of widely dispersed clonal lineages (Li et al. 2019; Guo et al. 2022; Tsushima et al. 2022; Lewis et al. 2024; Spanner et al. 2026a), such as the Pgt21 lineage (also known as Clade IX) found in Africa, Australia and Europe, and the Ug99 lineage (Clade I) now detected across Africa, the Middle East and Asia. Advances in sequencing technologies and genome assembly have allowed the generation of haplotype-phased genomes for a few *Pgt* lineages (Li et al. 2019; Spanner et al. 2026a), in which both haploid nuclear genomes are resolved. Such resources were not only instrumental in revealing the link between somatic hybridisation and the origin of Ug99, but have since provided further evidence of nuclear exchange events involved in the origin of other lineages of *Pgt* and epidemiologically important strains in other rust diseases such as wheat leaf rust (Sperschneider et al. 2023), oat crown rust (Henningsen et al. 2024), barley leaf rust (Spanner et al. 2026b) and wheat stripe rust (Holden et al. 2025; Hovmøller et al. 2025).

South America is a major wheat-growing area, producing between 5 and 10% of the world’s wheat harvest over the last 20 years (FAO 2025). The bulk of production occurs in the Southern Cone region including northeastern Argentina, (∼60% on average), southern Brazil (∼24%) and Uruguay (∼4%) (Garcia-Alonso et al. 2014; FAO 2025). Although not resulting in major epidemics recently, wheat stem rust is regularly detected in these regions, with yearly surveys conducted to track *Pgt* races by INIA in Uruguay and INTA in Argentina. However, South American isolates have not been well represented in genomic studies of *Pgt* conducted to date, so the population genetic structure in these regions and the relationship of local isolates to global lineages remain unknown.

Here, we present an analysis of *Pgt* samples collected over the 2020 and 2021 growing seasons in the wheat-growing regions of Argentina and Uruguay. A total of 91 samples of *Pgt* were collected from stem rust-infected wheat (n=69), barley (n=17), rye (n=4) and triticale (n=1) across seven Uruguayan districts (Paysandú, Rio Negro, Soriano, Colonia, San José, Flores, and Durazno, 78 samples) and four provinces in Argentina (Córdoba, Santa Fe, Entre Ríos and Buenos Aires, 13 samples) (**Fig.1, Table S1**). Single pustule isolates were purified on the susceptible wheat cultivar Morocco for subsequent race and genomic analyses using previously described procedures (Li et al. 2019).

A total of twelve races with different virulence profiles were identified from 57 isolates from Argentina and Uruguay (**Table 1**) after inoculation on seedlings of the North American differential set representing 20 wheat *Sr* resistance gene (*Sr5, Sr21, Sr9e, Sr7b, Sr11, Sr6, Sr8a, Sr9g, Sr36, Sr9b, Sr30, Sr17, Sr9a, Sr9d, Sr10, SrTmp, Sr24, Sr31, Sr38, SrMcN*) (Jin et al. 2008). Most of the Uruguayan isolates (36 out of 44) had one of two race types, RCCNC and RCCPC, that differ only by virulence on *SrTmp*, while an additional isolate had the related race type RCHPC (differs from RCCPC on *Sr9b*). The thirteen Argentinian isolates all showed related pathotypes (RCBNC, RHBNC, RHCNC, QCCNC and QHBNC) that differ from the common Uruguayan races and each other by virulence on only one or two differentials. This is consistent with single-step virulence mutations separating these races. Additional races identified in Uruguay had more divergent pathotypes including: QHHSF (3 isolates), QRFTC (1), QNKSC (1) and RHTTF (2). The latter race was the most broadly virulent, with virulence on 15 of 20 tested *Sr* genes. All isolates were virulent on *Sr5, Sr21, Sr9a, Sr10* and *SrMcN* while no isolates were virulent on *Sr9e, Sr24* and *Sr31*. None of the isolates collected in Argentina were virulent on *Sr11, Sr8a, Sr9b, Sr30, Sr9d, SrTmp* or *Sr38*, while virulence towards these genes was identified in Uruguayan isolates.

**Table 1.** Pathotyping results for 57 isolates on the North American Stem Rust differential set.

| **Isolate** | **Country** | **Year** | **Race code** | **Host** |
| --- | --- | --- | --- | --- |
| Pgt 20-134(2) | Argentina | 2020 | RHBNC | Wheat |
| Pgt 20-137(1) | Argentina | 2020 | RHBNC | Wheat |
| Pgt 20-18(1) | Argentina | 2020 | QHBNC | Wheat |
| Pgt 20-43(3) | Argentina | 2020 | RCBNC | Wheat |
| Pgt 20-44(1) | Argentina | 2020 | RCBNC | Wheat |
| Pgt20-11 | Argentina | 2020 | RHCNC | Wheat |
| Pgt20-13_is2 | Argentina | 2020 | RHBNC | Wheat |
| Pgt20-13_is5 | Argentina | 2020 | QCCNC | Wheat |
| Pgt20-136 | Argentina | 2020 | QCCNC | Wheat |
| Pgt20-142 | Argentina | 2020 | RHCNC | Wheat |
| Pgt20-143 | Argentina | 2020 | QCCNC | Wheat |
| Pgt20-20 | Argentina | 2020 | RHBNC | Wheat |
| Pgt20-35 | Argentina | 2020 | RHBNC | Wheat |
| DS10_21-0032 | Uruguay | 2020 | RCCPC | Wheat |
| DS10_21-0033 | Uruguay | 2020 | RCCPC | Wheat |
| DS10_21-0034 | Uruguay | 2020 | RCCNC | Wheat |
| DS10_21-0036 | Uruguay | 2020 | QNKSC | Wheat |
| DS10_21-0037 | Uruguay | 2020 | RCCNC | Wheat |
| DS10_21-0044 | Uruguay | 2020 | RCCNC | Wheat |
| DS10_21-0045 | Uruguay | 2020 | RCCNC | Wheat |
| DS10_21-0054 | Uruguay | 2020 | RCCPC | Wheat |
| DS10_21-0062 | Uruguay | 2020 | RCCPC | Barley |
| DS10_21-0065 | Uruguay | 2020 | RCCNC | Barley |
| DS10_21-0068 | Uruguay | 2020 | QRFTC | Wheat |
| DS10_21-0073 | Uruguay | 2020 | RCCNC | Wheat |
| DS10_21-0076 | Uruguay | 2020 | RCCNC | Wheat |
| DS10_21-0077 | Uruguay | 2020 | RCCNC | Wheat |
| DS10_21-0079 | Uruguay | 2020 | RCCNC | Wheat |
| DS10_21-0080 | Uruguay | 2020 | RCCNC | Wheat |
| DS10_21-0083 | Uruguay | 2020 | RCCNC | Wheat |
| DS10_21-0086 | Uruguay | 2020 | RHTTF | Wheat |
| DS10_21-0089 | Uruguay | 2020 | RCCNC | Wheat |
| DS10_21-0091 | Uruguay | 2020 | RCCNC | Wheat |
| DS10_21-0092 | Uruguay | 2020 | RCCNC | Barley |
| U3674 | Uruguay | 2021 | RHTTF | Wheat |
| U3675 | Uruguay | 2021 | QHHSF | Wheat |
| U3676 | Uruguay | 2021 | RCCPC | Wheat |
| U3677 | Uruguay | 2021 | RCHPC | Wheat |
| U3680 | Uruguay | 2021 | RCCNC | Wheat |
| U3681 | Uruguay | 2021 | QHHSF | Wheat |
| U3758 | Uruguay | 2021 | RCCPC | Wheat |
| U3761 | Uruguay | 2021 | RCCPC | Wheat |
| U3764 | Uruguay | 2021 | RCCPC | Barley |
| U3765 | Uruguay | 2021 | RCCNC | Wheat |
| U3767 | Uruguay | 2021 | RCCNC | Wheat |
| U3773 | Uruguay | 2021 | RCCPC | Triticale |
| U3780 | Uruguay | 2021 | QHHSF | Wheat |
| U3789 | Uruguay | 2021 | RCCPC | Barley |
| U3798 | Uruguay | 2021 | RCCNC | Barley |
| U3799 | Uruguay | 2021 | RCCPC | Barley |
| U3801 | Uruguay | 2021 | RCCNC | Barley |
| U3808 | Uruguay | 2021 | RCCPC | Wheat |
| U3810 | Uruguay | 2021 | RCCNC | Barley |
| U3811 | Uruguay | 2021 | RCCNC | Wheat |
| U3813 | Uruguay | 2021 | RCCNC | Wheat |
| U3815 | Uruguay | 2021 | RCCNC | Wheat |
| U3816 | Uruguay | 2021 | RCCPC | Barley |

To explore genetic diversity in this *Pgt* population, we generated whole-genome or RNA transcriptome sequence data from 86 isolates. RNA was extracted using the Qiagen Plant RNeasy kit (Cat No: 74904) from 78 field samples from Uruguay that had been stored in RNAlater (Thermo Fisher Scientific, Waltham, MA, USA) and was sequenced by Azenta (Burlington, MA, USA) using 150 bp paired-end read libraries and the Illumina NovaSeq 6000 platform. Genomic DNA was extracted from eight isolates from Argentina using the Omniprep^TM^ Genomic DNA isolation kit (#786-136, G-Biosciences) and sequenced by the University of Minnesota Genomics Center (UMGC, St. Paul, MN) using 150 bp paired-end Nextera Flex library preparation and the Illumina NovaSeq 6000 platform. The sequence data for these samples was combined with that of 150 other global *Pgt* isolates published previously (**Table S1**) (Duplessis et al. 2011; Upadhyaya et al. 2015; Chen et al. 2017; Rutter et al. 2017; Salcedo et al. 2017; Lewis et al. 2018; Li et al. 2019; Kangara et al. 2020; Guo et al. 2022; Tsushima et al. 2022). Trimmed reads were aligned to the Pgt21-0 reference genome (Li et al. 2019) with the called SNPs used to generate a Maximum Likelihood (ML) tree as described previously (Spanner et al. 2026a). Nine samples were excluded because allele frequency analysis indicated a mixture of genotypes of two or more *Pgt* isolates (**Fig. S2**). The remaining 77 South American isolates fell into three different clonal groups in the ML tree (**Fig. 2**). The majority (69 isolates, ∼90%) belong to a single clonal lineage (**Fig. 2**, clade shown in blue) that is unrelated to any of the previously sequenced *Pgt* isolates or lineages in the tree. This group includes the 8 Argentinian isolates and 61 of the Uruguayan isolates and encompasses the two most common race types (RCCPC and RCCNC) as well as the closely related pathotypes RCHPC, RCBNC, RHBNC, RHCNC, QCCNC and QHBNC. This confirms that these isolates are related members of a clonal lineage that has evolved by single-step mutations to overcome individual resistance genes. Four additional isolates from Uruguay (U3768, U3770, U3772 and U3769) form another unique clonal lineage (**Fig. 2**, clade shown in yellow) divergent from other sequenced global isolates/lineages of *Pgt*. None of these isolates were pathotyped in this study so the race types represented by this novel South American lineage are unknown. Interestingly, U3769, U3770 and U3772 were all collected from rye, while the next most closely related isolates in the tree (DE-04, DE-05, DE-06 and DE-07) were also collected from rye (**Fig. 2, Table S1**) and have been previously characterized as *P. graminis* f. sp. *secalis* (*Pgs*) (Bryan et al. 2024). This could indicate that the Uruguayan isolates are also *Pgs*. Finally, another four isolates from Uruguay (DS10_21-0032, DS10_21-0086, U3674 and U3675) formed a clonal group that includes isolates from Israel, Egypt, Kenya and Madagascar, collected between the 1980s and the mid-2000s (**Fig. 2**, clade shown in red, **Fig. S1**) and representing a branch of the previously characterized clade III (Guo et al. 2022; Lewis et al. 2024; Spanner et al. 2026a). The *Pgt* isolates from South America included in this clade comprised two races specific to the group, RHTTF (two isolates) and QHHSF (one isolate), which are similar to pathotypes of the previously characterised isolates in this lineage (87MDG1054A, 84KEN54B, 99EGY5B: RRTTF) (Guo et al. 2022). The fourth South American isolate placed in this group was assigned the race type RCCPC, typical of the larger South American *Pgt* lineage, and may therefore be derived from a mixed sample. A SplitsTree network (Huson and Bryant 2024) of 80 South American *P. graminis* f. sp. *tritici* isolates generated from 202,603 SNPs across the entire Pgt21-0 genome supports a lack of sexual recombination in these populations (**Fig. S1**).

**Fig. 1.**
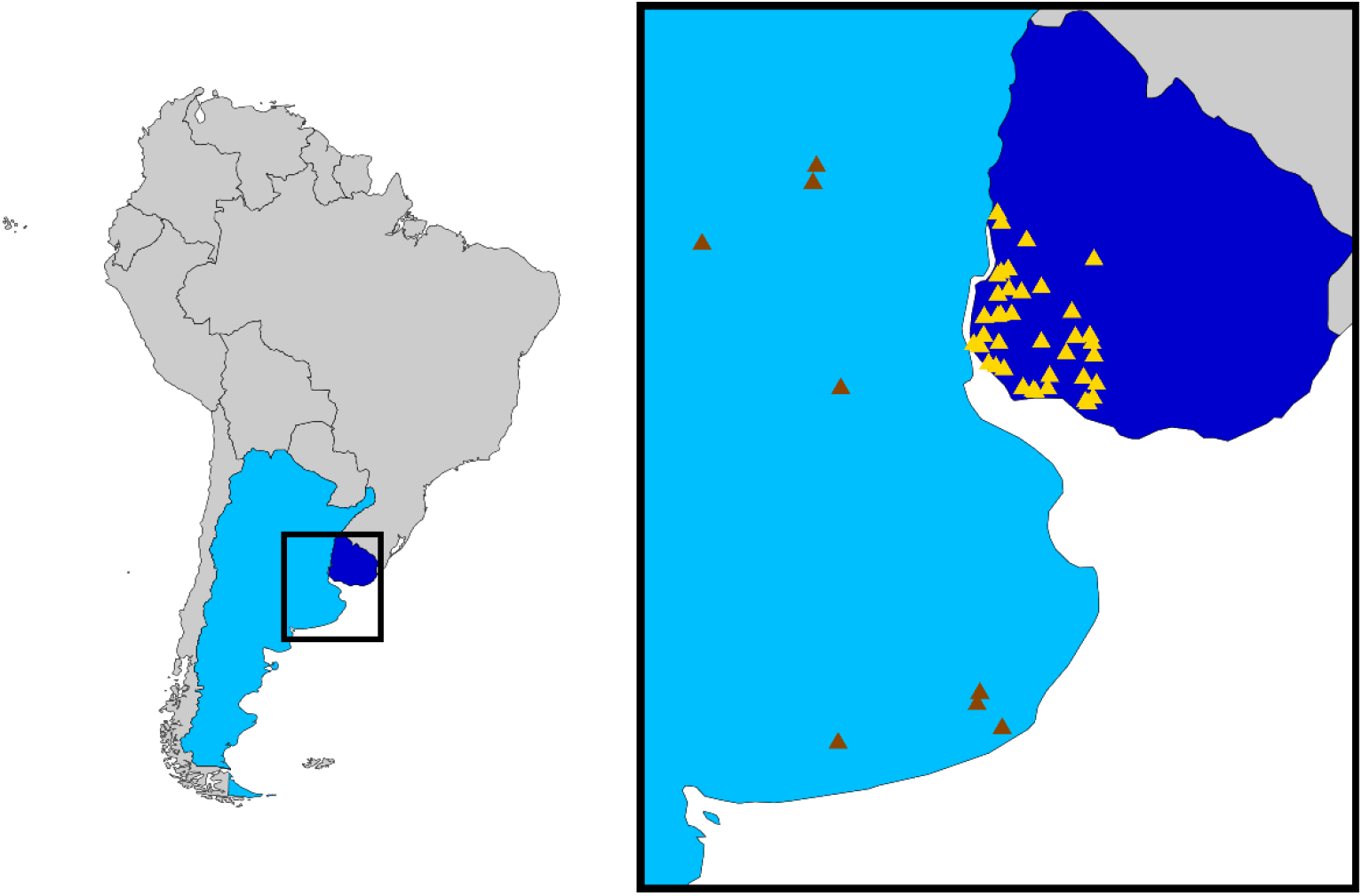
Map of South America showing the sites (brown and yellow triangles) in Argentina (light blue) and Uruguay (dark blue) where *P. graminis* f. sp. *tritici* samples used in this study were collected.

**Fig. 2.**
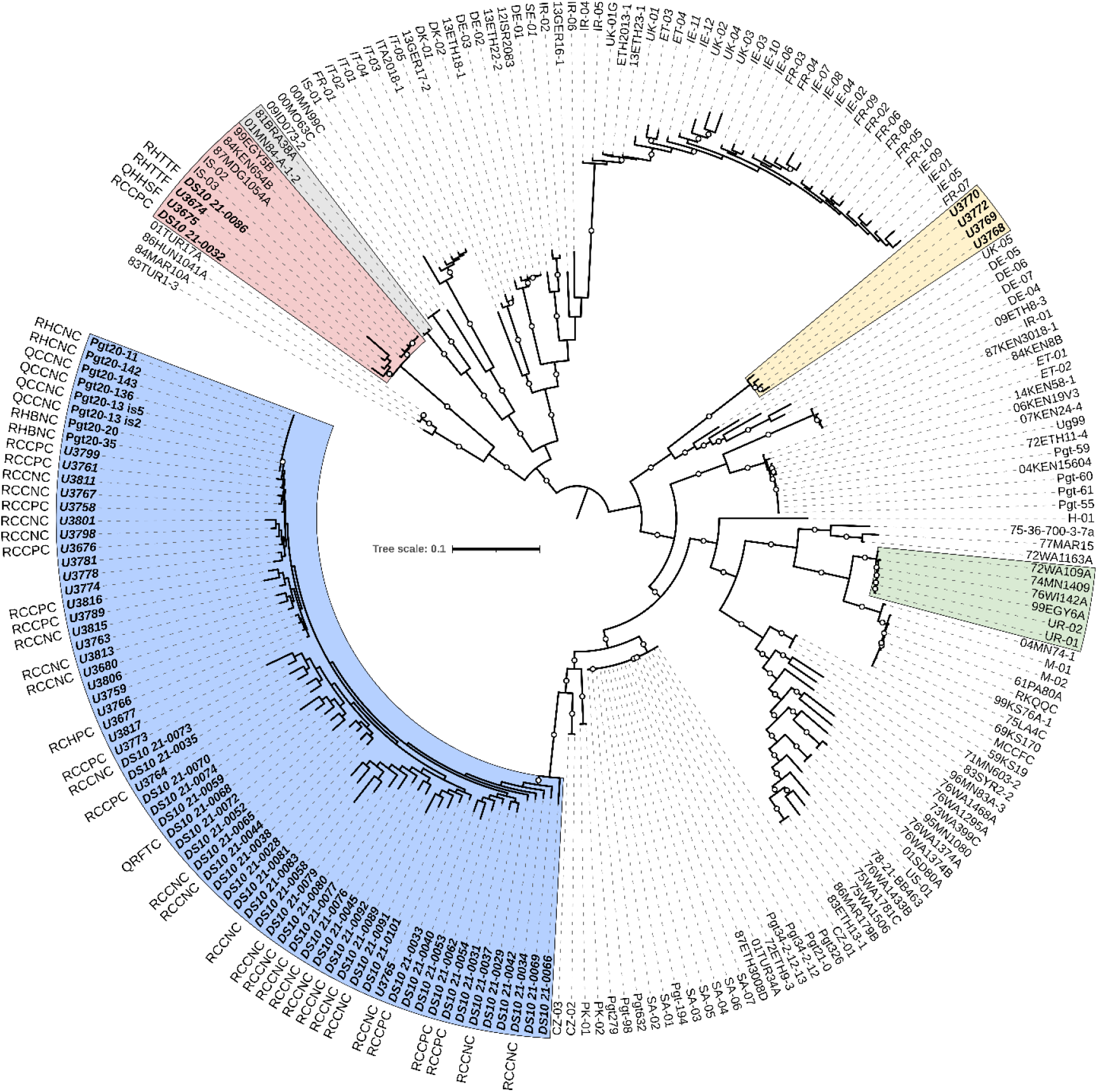
Maximum likelihood phylogenetic tree of 225 *Puccinia graminis* f. sp. *tritici* isolates based on SNPs called against the complete Pgt21-0 genome (202,603 SNPs). Bootstrap values over 80% are shown as circles at branch midpoints. Scale is the mean substitutions per site. Bold text indicates *Pgt* isolates from this study, with race codes shown for pathotyped isolates. Font in italics indicates isolates with RNAseq data, while other isolates are based on whole genome sequence data. Blue and yellow boxes represent novel lineages identified in this study, while grey, green, and red boxes indicate known clades containing isolates from South America (respectively: 81BRA38A from Guo et al. 2022; UR-01 and UR-02 from Lewis et al. 2018; DS10 21-0086, U3674, U3675 and DS10 21-0032 from this study).

For the eight Argentinian isolates in the predominant South American lineage with whole genome sequence data, we also performed a *k*-mer-containment analysis against the nuclear haplotypes hap01 to hap07 that have been defined in whole genome references for Pgt21-0 (clade IX), Ug99 (Clade I), TTRTF (Clade III-B) and TKTTF (Clade IV-B) (Spanner et al. 2026a). Genomic reads were screened with mash v2.3 sketch (-s 100000) (Ondov et al. 2019), which showed that none of these known haplotypes are contained by *Pgt* isolates in this lineage (**Table S2**). Therefore, we conclude that they are not related to any of these global lineages by nuclear exchange.

Our analysis also confirmed previous reports that a South American isolate collected in Brazil in 1981 (81BRA38A) grouped with 01MN84-A-1-2, an isolate from the USA collected in 2001 (Guo et al. 2022) (**Fig. 2**, shown in grey), likely illustrating a lineage that has been widely distributed throughout the American continent. Savva et al. (2025) generated a phylogenetic tree that grouped Uruguayan isolates UR-01 and UR-02 collected in 2011 (Lewis et al. 2018) with isolates collected in the USA in the 1970s, including the US isolate 74MN1409, collected in 1974 (**Fig. 2**, shown in green). Our analyses placed Egyptian isolate 99EGY6A collected in 1999 into this group, a result that corroborates the previous grouping of this isolate with 74MN1409 (Guo et al. 2022). Thus, this group represents a long-lived and globally dispersed clonal lineage.

These findings strongly suggest spore dispersal over large geographic distances manifesting as gene flow between Eurasia, Africa and South America on the one hand and between North and South America on the other. In the case of stem rust and rust diseases in general, due to their potential for global spread, breeding programs must include not only responses to the local landscape of fungal populations, but also the potential for incursions from elsewhere. Our results support the need for further investments to genotypically characterise larger *Pgt* collections from South America and the incorporation of these genotype data into the existing global information network.

In summary, we surveyed a specific wheat-growing region spanning Northeast Argentina and Southwest Uruguay with isolates collected over two years. The results indicated that the current South American *Pgt* population includes three clonal lineages, two of which are unique to the region and not represented in samples from other regions of the world, while the third represents a globally dispersed lineage. Historical samples from South America represent two additional international lineages, but it is not known if these can still be detected in the region. Importantly, there was no evidence of sexual recombination giving rise to new genetic diversity in these populations despite the presence of barberry in Brazil and nearby regions (Rezaei and

Sarkhosh 2025). The genotypic information gathered from this collection of South American *Pgt* isolates and their pathotypes can be harnessed to expand the existing *Pgt* pangenome effort (Spanner et al. 2026a). With such resources in hand, we will be well-positioned to detect *Avr* allelic variants to increase the *Pgt Avr* gene atlas under construction (Spanner et al. 2026a). This data, in turn, can be used in high-throughput surveillance platforms such as MARPLE (Savva et al. 2025), contributing to a robust and sensitive monitoring system.

---

**Fig. S1.**
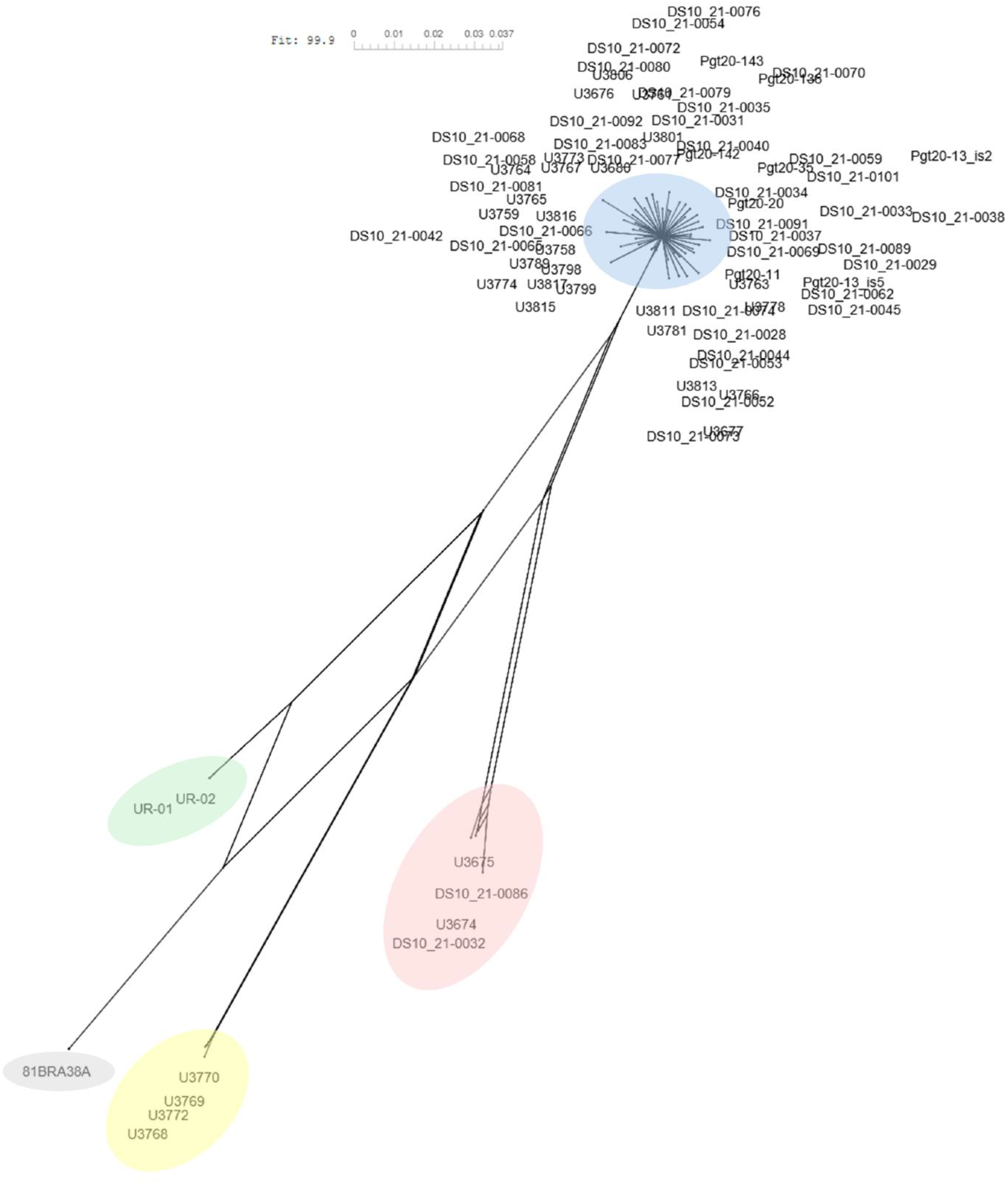
SplitsTree network of 80 South American *P. graminis* f. sp. *tritici* isolates generated from 202,603 SNPs across the entire Pgt21-0 genome. Heterozygous sites were converted to missing. Tree scale is mean substitutions per site. Blue and yellow represent novel lineages identified in this study, while grey, green, and red indicate known clades containing isolates from South America (respectively: 81BRA38A from Guo et al. 2022; UR-01 and UR-02 from Lewis et al. 2018; DS10 21-0086, U3674, U3675 and DS10 21-0032 from this study).

**Fig. S2.**
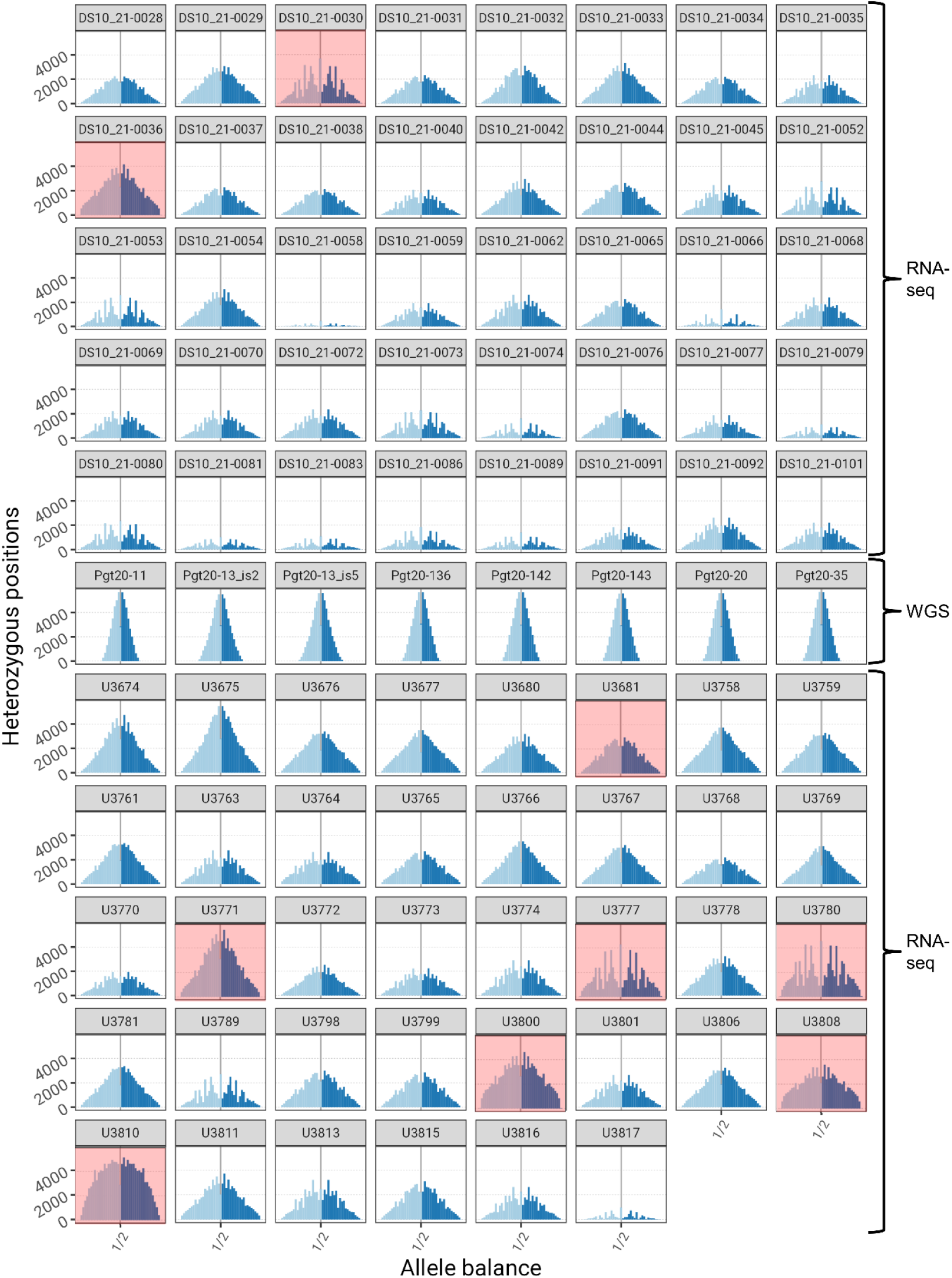
Allele frequency distribution of 341,396 heterozygous biallelic sites in the 86 South American *P. graminis* f. sp. *tritici* isolates sequenced in this study. Nine isolates were excluded from further analyses for showing signs of contamination (highlighted in red).

